# *De novo* motor learning creates structure in neural activity space that shapes adaptation

**DOI:** 10.1101/2023.05.23.541925

**Authors:** Joanna C. Chang, Matthew G. Perich, Lee E. Miller, Juan A. Gallego, Claudia Clopath

## Abstract

Animals can quickly adapt learned movements in response to external perturbations. Motor adaptation is likely influenced by an animal’s existing movement repertoire, but the nature of this influence is unclear. Long-term learning causes lasting changes in neural connectivity which determine the activity patterns that can be produced. Here, we sought to understand how a neural population’s activity repertoire, acquired through long-term learning, affects short-term adaptation by modeling motor cortical neural population dynamics during *de novo* learning and subsequent adaptation using recurrent neural networks. We trained these networks on different motor repertoires comprising varying numbers of movements. Networks with multiple movements had more constrained and robust dynamics, which were associated with more defined neural ‘structure’—organization created by the neural population activity patterns corresponding to each movement. This structure facilitated adaptation, but only when small changes in motor output were required, and when the structure of the network inputs, the neural activity space, and the perturbation were congruent. These results highlight trade-offs in skill acquisition and demonstrate how prior experience and external cues during learning can shape the geometrical properties of neural population activity as well as subsequent adaptation.

## Introduction

From walking to grasping objects, movement enables us to interact with the world. Mastering a skill requires many hours of practice, be it during development or in adulthood. In contrast to long-term skill learning, adapting existing skills to environmental perturbations is a much faster process: after learning to ride a bike, adapting to foggy weather or uneven roads is much easier. Existing motor repertoires acquired through long-term learning likely form the foundation for short-term motor adaptation, but it is unclear how different repertoires can affect adaptation, even for common experimental perturbations like visuomotor rotations (VR)^1^ or force fields ^2^. This lack of understanding stems from the experimental challenge of characterising an animal’s entire behavioural repertoire learned throughout their lifetime, and implies that the interplay between available neural activity patterns and rapid adaptation remains largely unknown ^3^.

Recent work has focused on the coordinated activity of neural populations to begin to shed light on the neural basis of motor adaptation ^4–6^. In this neural population view, brain function is not build upon the independent activity of single neurons, but rather on specific patterns of neural co-variation (from now on, simply ‘activity patterns’) ^7–9^. In practice, these activity patterns can be examined by building a neural (state) space where each point denotes the state of the neural population. Numerous studies (e.g. Refs. 10–15) have found that the activity of even hundreds of simultaneously recorded neurons is well captured by relatively few population-wide activity patterns, an observation consistent with neural population activity being constrained to a low-dimensional surface—a ‘neural manifold’—that can be estimated by applying a dimensionality reduction method ^16–18^. Interestingly, this neural manifold is likely shaped, at least partly, by the underlying connectivity of the network ^15, 19–21^. Previous work suggests that long-term learning causes changes in circuit connectivity ^22–25^, which may in turn change the geometrical properties of an existing neural manifold as well as the activity within it — the so-called ‘latent dynamics’. In contrast, short-term motor adaptation may be achieved without significant changes in circuit constraints ^6, 26^. Indeed, using a brain-computer interface that mapped neural activity onto computer cursor movements, Sadtler et al. ^15^ showed that it is easier to adapt to perturbations that require only neural states that lie within the existing neural manifold: while animals can learn to produce activity patterns within the existing manifold in a matter of minutes or hours, producing activity patterns outside the existing manifold takes several days ^15, 19^. To learn outside-manifold perturbations, new activity patterns need to be used, which may necessitate changing the synaptic connectivity of the circuit ^19^. Combined, these results suggest that long-term motor learning changes the circuit connectivity to create new neural population latent dynamics whereas short-term motor adaptation may reuse existing ones ^27^.

Yet, studying the interplay between an animal’s existing motor repertoire and its ability to adapt a known behaviour remains challenging due to the complex organisation of existing motor skills within neural activity space. Similar behaviors (e.g., various wrist manipulations or grasping tasks) may share similarly oriented task-specific neural manifolds ^12^, while dissimilar behaviors (e.g., reaching and walking in mice) may have almost orthogonal manifolds even if they share similar movements ^14^. Furthermore, experimentally, it is impossible to quantify the entire repertoire of motor skills that an animal has, which poses a challenge to investigate how its behavioral repertoire influences motor adaptation. Moreover, in virtually all motor adaptation studies, animals must adjust to perturbations on a specific laboratory task they have already learned, and adaptation is only examined with respect to a ‘baseline period’ (e.g. Refs. 1,2,4– 6,15); this approach ignores, for practical reasons, the relationship between the adapted behavior and all the other motor skills the animal has previously acquired.

To overcome the experimental challenges of assessing an animal’s lifelong experience, here we examined how the existing motor repertoire can affect adaptation differently using recurrent neural networks (RNNs). Similar networks have been able to reproduce motor output and key features of latent dynamics from experimental recordings ^28–31^, including during adaptation ^26, 32^. Here, we modeled the latent dynamics of the motor cortex during *de novo* learning and subsequent adaptation. We trained our networks on different repertoires with varying numbers of movements, using movement trajectories modified from experimental recordings of monkey reaches ^6, 33^. We hypothesized that networks with larger motor repertoires would adapt to perturbations more easily since they are already able to produce a broader set of activity patterns. In conjunction, we investigated how both the organization of latent dynamics in neural space and the structure of the cues present during learning impacted adaptation.

By systematically training networks in two stages comprising of *de novo* learning and subsequent adaptation, we found that larger repertoire networks could adapt to perturbations more quickly, but only under certain circumstances. The way the latent dynamics of multiple movements were organized in neural space shaped subsequent adaptation: adaptation was facilitated when the latent dynamics were organised in a way that was congruent with the motor output changes required by the perturbation, and when only small changes in motor output were needed. This suggests that ease of adaptation is affected not only by its relation to the existing manifold ^15, 19^, but also by the organization of the latent dynamics within it, and this organization is affected by past learning experiences. This observation also highlights an inherent trade-off in skill-acquisition: mastery of more movements better defines the structure of the neural manifold. This, in turn, facilitates adaptation that requires small changes in behavior, but potentially harms adaptation that requires large changes or when the learning experience is different.

## Results

### Probing the impact of *de novo* learning on subsequent adaptation with RNNs

To understand how a neural population’s existing activity patterns affect its ability to change its activity, we used RNNs to model motor cortical neural population dynamics following *de novo* learning and subsequent adaptation (Figure 1A).

**Figure 1:**
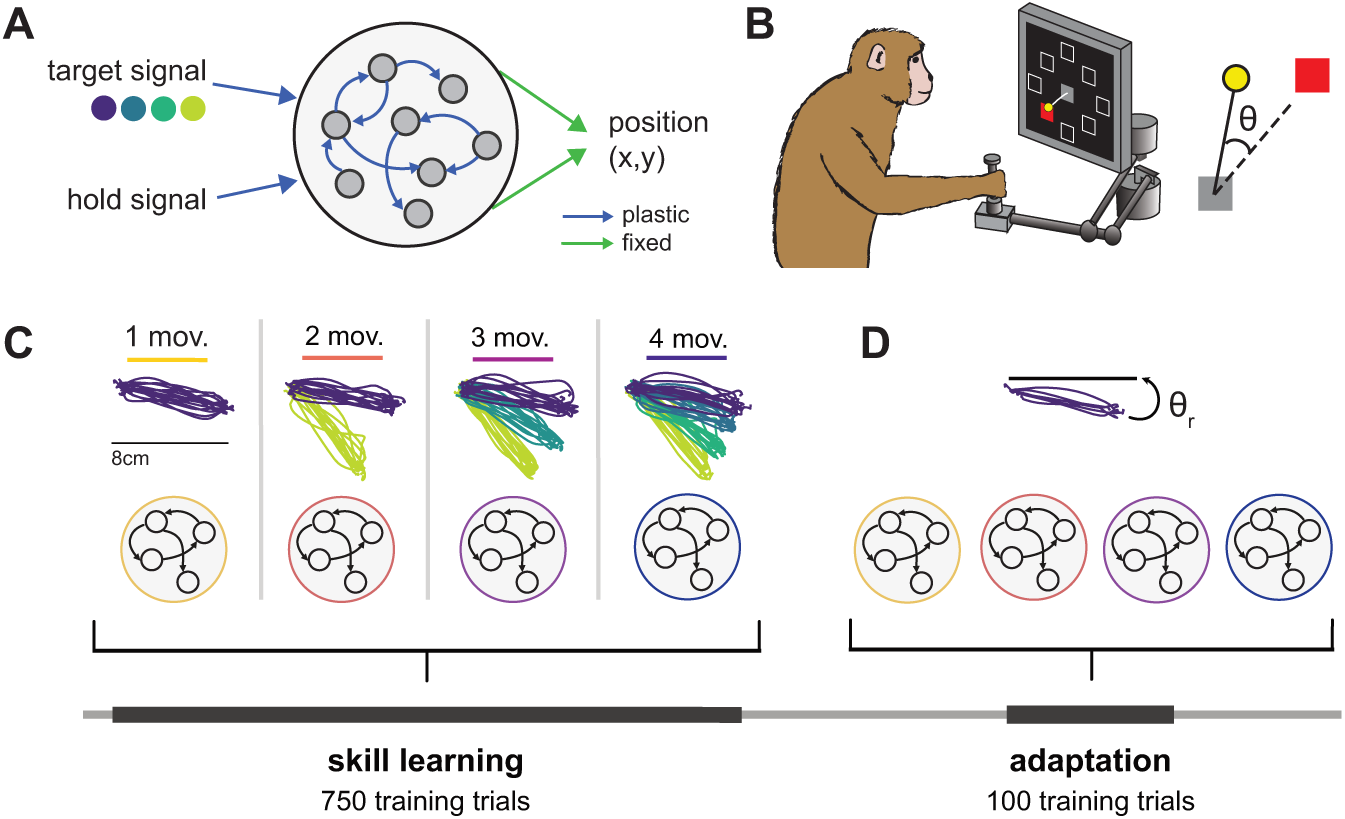
Probing *de novo* learning and adaptation with RNNs. **A**. RNNs were trained to produce ‘hand positions’ as output. They were given a target signal that specifies the reach direction, and a hold signal that indicated movement initiation. The target signal indicated the angular direction, unless otherwise specified. The input and recurrent weights were learned (plastic) while the output weights were fixed, unless otherwise specified. **B**. RNNs were trained on hand trajectories modified from experimental recordings from monkeys performing a centerout reaching task (left). Subsequent adaptation was studied using a classic visuomotor rotation paradigm (VR, right), in which visual feedback is rotated by a fixed angle around the center of the workspace. **C**. Networks were trained on repertoires with different numbers of movements (from one to four) to model *de novo* learning. All multi-movement networks (2 mov., 3 mov. and 4 mov.) covered the same angular range. **D**. Networks were later trained to counteract VR perturbations for only the one common movement to understand the influence of the existing motor repertoires on adaptation.

To model *de novo* skill learning, we trained the networks on repertoires with different numbers of movement directions. The movements were modified from experimentally recorded center-out reaches from monkeys (data from Perich et al. ^6^, Figure 1B, Methods). For each reach, monkeys were first presented with a visual target; after a variable delay period, a go cue indicated they could execute the movement. To address our hypothesis that having a larger motor repertoire would facilitate adaptation, we used repertoires of different sizes, ranging from one to four movement directions (Figure 1C). Single movement repertoires comprised a single reach to −10° while multi-movement repertoires comprised movements in directions equally spaced between −10° and −50°, unless otherwise specified. Note that we intentionally used networks with only up to four movements to make our simulation results intelligible by comparing simple models. Importantly, all repertoires included the −10° direction, so all networks learned to produce this movement, allowing for comparisons across networks trained on different repertoires. Networks were given the angular direction of these movements as inputs along with a ‘go’ cue (occurring 0.5–1.5 s after the direction cue) to mimic the instructed delay reaching task performed by the monkeys (Methods).

To model motor adaptation, we subsequently trained these networks to counteract a visuomotor rotation (VR) on the shared movement direction they had all learned (Figure 1D). VR is a commonly used experimental paradigm to examine motor adaptation ^1, 34–36^ in which a rotational transformation is applied to the motor output. In this case, we applied a 10° rotation (counterclockwise), which the networks had to counteract by producing output in the opposite direction. By probing adaptation on only one common movement, we could assess how entire motor repertoires contribute to adaptation of a given movement, and compare the adaptation performance across repertoires.

### Multiple movements produce more constrained and robust dynamics that are structured in neural activity space

We first trained networks to produce repertoires comprising one to four different movements to understand the resulting differences in underlying network activity. Following the initial *de novo* learning phase, all the networks were able to learn each of the repertoires (Figure 2A) with comparable performance (Figure 2B), as quantified by the mean squared error between target movement trajectories and produced movement trajectories. This indicates that any differences in their ability to adapt will not be due to how accurately they can generate motor output, but rather to differences in the network dynamics that can be produced based on their acquired motor repertoires.

**Figure 2:**
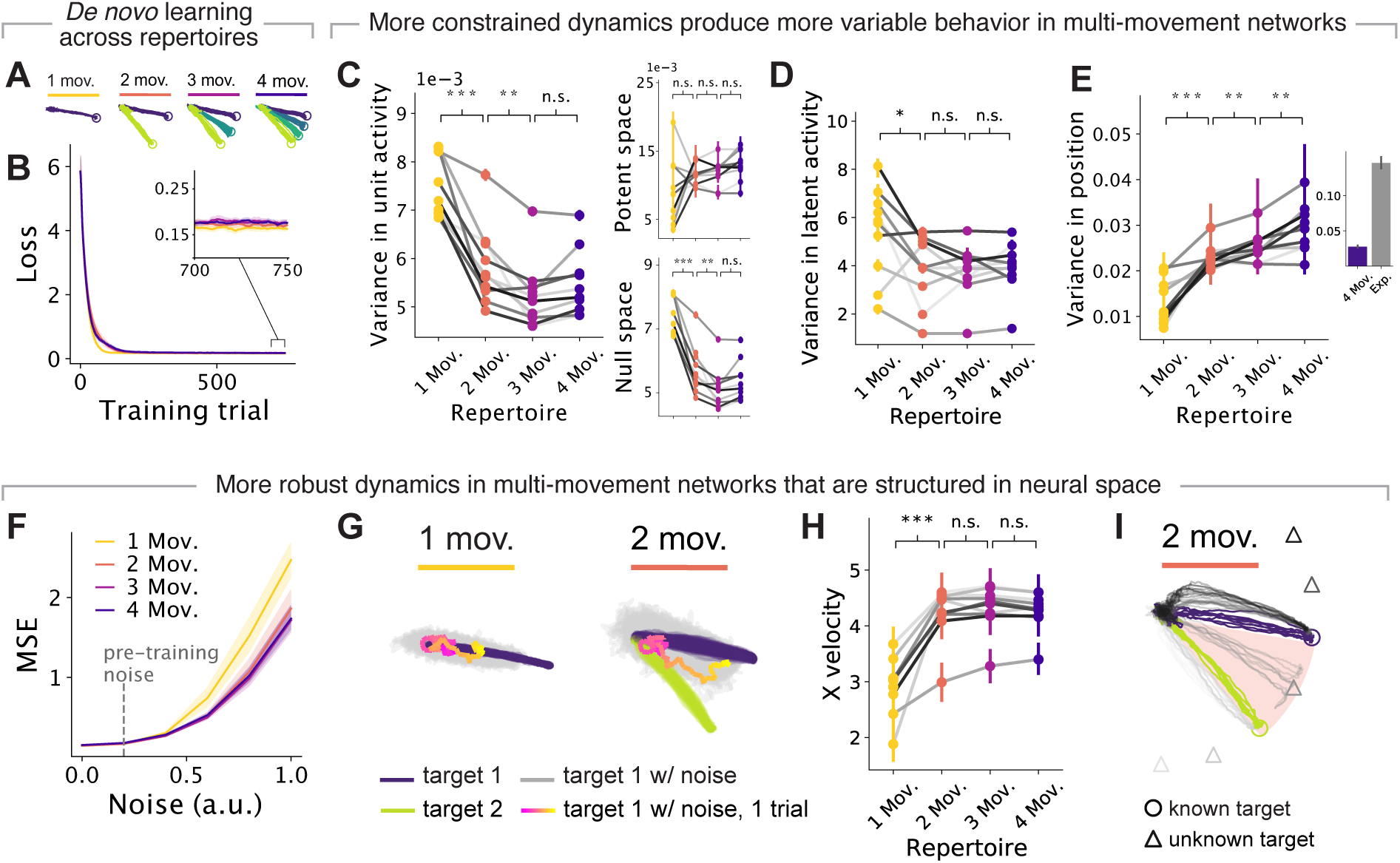
Networks that have acquired multiple movements produce more constrained and robust dynamics that are structured in neural activity space. **A**. Motor output of networks trained on different repertoires following *de novo* learning. **B**. Loss during motor skill training. Loss was calculated as the mean-squared error between the network output and target positions. Line and shaded surfaces, smoothed mean and 95% confidence interval across networks of different seeds. **C-F, H**. Following *de novo* learning, networks were tested on the one shared movement. **C**. Variance in unit activity for networks trained on different repertories. Inset: Variance in the output-null subspace and output-potent subspace of the unit activity with respect to the produced output. Individual lines, different random seeds; Circles and error bars, median and 95% confidence intervals with bootstrapping. *** denotes 0.001, ** 0.01, * 0.05 for Wilcoxon signed rank tests. **D**. Same as Panel C, but for the variance in latent dynamics. **E**. Same as Panel C but for variance in motor output. Inset: variance in position in a single reach for networks trained on 4 movements to that for monkeys trained on the center-out reach task (data from Ref. 6). Note that the networks, which only know a few movements, generally have less variance than monkeys that know many more movements. **F**. Mean-squared error of output when noise of increasing magnitude is added to the neural activity. Line and shaded area, median and 95% confidence interval. **G**. Motor output with (grey, pink gradient) and without (purple, green) increased noise added (*η* = 1) for example networks with one- and two-movement repertoires. Color gradient, time-course of the movement execution (legend). **H**. Same as Panels C–E but for velocity in the *x* direction for motor output when noise (*η* = 1) is added as per Panel G. **I**. Motor output for different target cues for a sample network with two-movement repertoire. Colors, different target cues (target cues that network has been previously trained on have higher opacity); circles, target endpoints for each movement; pink background, range of known movements.

While performance was similar, we had predicted that differences in the motor repertoire will lead to differences in the neural dynamics produced by the networks. To compare across different repertoires, we examined the dynamics as the networks were producing the movement that was shared across all repertoires (−10°).

We focused on network activity during both preparation and execution of the same target motor output. We calculated the population latent dynamics by projecting the activity of all 300 units in our networks onto a lower-dimensional neural manifold identified by performing Principal Component Analysis ^7, 37^ (PCA) (Methods). The main differences in the activity produced by networks with different repertoires was between single-movement and multi-movement (2, 3, or 4 movements) networks: multi-movement networks had less variance in both single unit activity (Figure 2C left, *P* = 9.8 *·* 10*^−^*^4^, Wilcoxon signed rank test; Figure S1A-C) and latent dynamics (Figure 2D, *P* = 0.02, Wilcoxon signed rank test; Figure S1D-F) than single-movement networks, suggesting that the network activity becomes more constrained when multiple movements need to be embedded in the neural space. Intriguingly, despite their lower variance in activity, multi-movement networks had greater variance in motor output (Figure 2E, *P* = 9.8 *·* 10*^−^*^4^, Wilcoxon signed rank test; Figure S1G-I; Figure S2)—which was more comparable to experimental reaches (Figure 2E right)—even though all the networks were performing the same movement. How can these apparently contradictory observations be reconciled? We hypothesized that the more variable latent dynamics of single-movement networks may lie on directions of activity space that do not affect motor output, i.e., those defining the ‘output-null’ subspace ^38^. Separately computing the variance of the latent dynamics within the output-null subspace and the ‘output-potent’ subspace (i.e., the dimensions that affect the network output), confirmed this prediction: the greater variability of the latent dynamics of single-movement networks was largely confined to the output-null subspace, and thus did not lead to more variable motor output (Figure 2C right, see Methods). These results also held when networks were trained on synthetic reaches that were more stereotyped than actual monkey reaches (Figure S3). Thus, greater constraints in the network dynamics were characteristic of multi-movement networks, but these more constrained dynamics did not necessarily translate into more consistent behavioural output.

**Figure 3:**
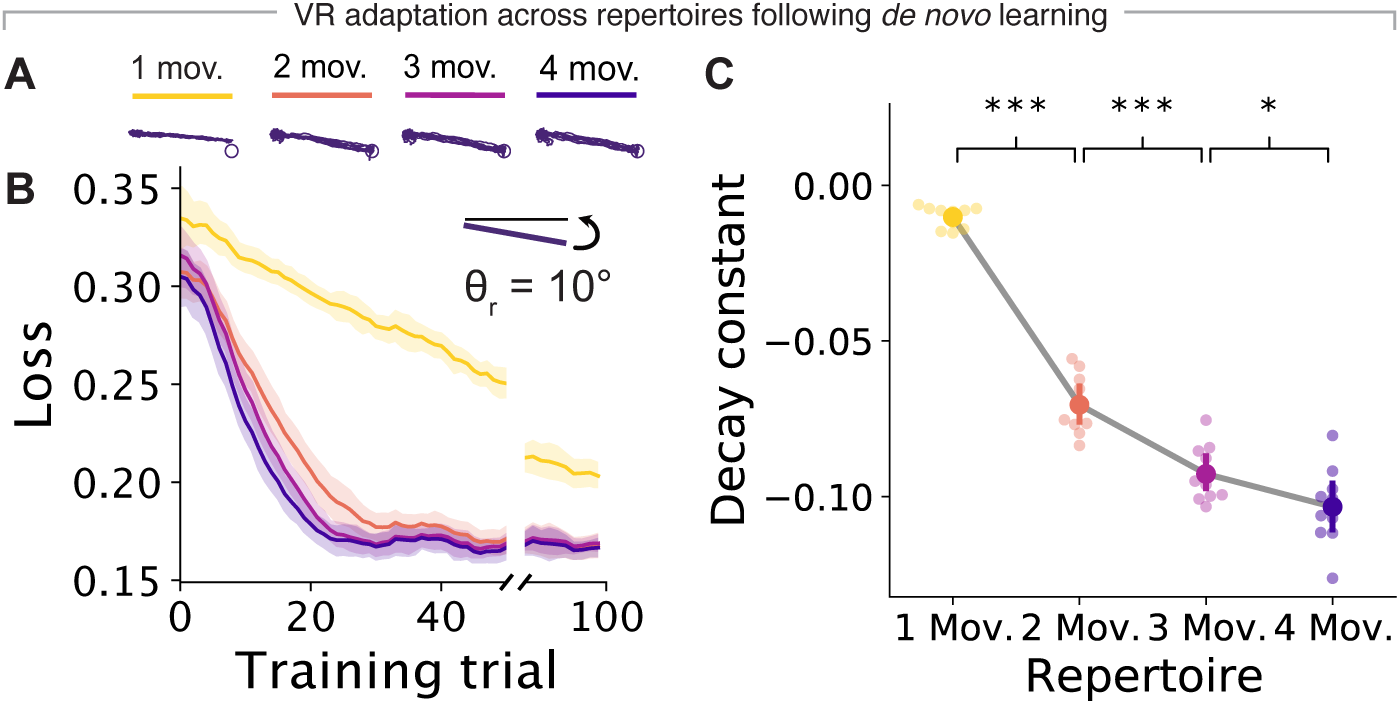
Networks with larger repertoires can adapt to perturbations more easily. **A**. Motor output of networks trained on different repertories (legend) following adaptation to a counterclockwise VR perturbation. **B**. Loss during adaptation, calculated as the mean-squared error between the network output and target positions. Line and shaded surfaces, smoothed mean and 95% confidence interval across networks of different seeds. **C**. Decay constants for exponential curves fitted to the loss curves in (**B**). Circles and error bars, means and 95% confidence intervals with bootstrapping. *** denotes 0.001, ** 0.01, * 0.05 for paired t-tests.

How may these constraints affect the network’s ability to generate robust latent dynamics driving motor output? We predicted that they may lead to greater robustness against noise ^39^. We examined this by changing the amount of simulated noise applied to the unit activity, and saw that multi-movement networks were indeed more robust against higher noise levels than single-movement networks (Figure 2F). To probe this further, we increased the noise five-fold compared to pre-training and measured how the output was affected (see examples in Figure 2G, with the two-movement network representative of multi-movement networks). Intriguingly, while motor output for single-movement networks circled back and became twisted, that for two-movement networks shifted toward previously learned outputs at this high noise level (Figure 2G). This systematic shift suggests that learning multiple movements creates structure in the neural activity space that maps to structure in the motor output, that is, the trajectories described by the latent dynamics driving each movement are organised in neural space in a way that is congruent with that of the movements. With this underlying ‘congruency’, noise in the activity space caused two-movement networks to explore other activity states that led to movements intermediate to those that had been previously learned (Figure 2G). This led to mostly linear trajectories with greater forward movement, as quantified by the velocity in the *x* direction during execution (Figure 2H, *P* = 9.8 *·* 10*^−^*^4^, Wilcoxon signed rank test), in contrast to the twisted and tangled movements produced by single-movement networks. Moreover, with this underlying structure, two-movement networks were able to generate intermediate movements that they had not previously learned when probed with the appropriate input signal (Figure 2I, Figure S4B,C). Combined, these results suggest that learning multiple movements creates structure in the neural activity space that effectively makes more intermediate activity patterns available. The availability of these additional activity patterns may facilitate adaptation.

### Networks with larger repertoires can adapt to a perturbation more easily

To directly assess the prediction that the additional structure in neural activity space of multi-movement networks may facilitate adaptation, we applied the same small VR perturbation of 10° to networks with different movements (Figure 3A,B). In general, networks that had learned larger repertoires were able to adapt more quickly (Figure 3B,C, Figure S4D,E), and without catastrophic forgetting (Figure S5). Comparison of adaptation performance across networks with different motor repertoires revealed two trends. First, all three types of multi-movement networks adapted much more rapidly than single-movement networks, as predicted. Within 100 adaptation training trials, multi-movement networks converged to a performance comparable to baseline. Single-movement networks, in contrast, could not fully adapt and had average errors 20% larger than that of the multi-movement networks (Figure 3B). Second, within multi-movement networks, those with larger repertoires also adapted more quickly than those with smaller repertoires (Figure 3C). These differences were smaller than those observed between single and multi-movement networks, suggesting that there may be two different processes at play. To understand the neural underpinnings of these two processes, we further examined the dynamics and behavior of the networks.

### The structure of neural activity in multi-movement networks is responsible for their patterns in adaptation

Following *de novo* learning, there were large differences in activity between single-movement and multi-movement networks, and subtle differences among multi-networks with different repertoires (Figure 2). These differences were reflected in the trends observed during adaptation (Figure 3). To understand the basis for these differences, we examined how activity is structured in neural space, since it was a differentiating characteristic for multi-movement networks. Single-movement networks lack multiple latent trajectories that can be structured with respect to one another, so we cannot directly manipulate this structure to assess its effects in single-movement networks. Instead, we focus on how structure affects adaptation in multi-movement networks.

We quantified structure in the neural space by examining the organization of the latent trajectories in multi-movement networks during preparation and execution. During both the preparatory and execution epochs, the latent trajectories were organized congruently to the movements themselves, with movements reaching to adjacent targets also adjacent in neural space (Figure 4A-C). This suggests that changes in the neural space may consequently lead to congruent changes in the motor output. To quantify this congruence, we measured the Euclidean distances in the neural manifold between the latent trajectories at corresponding time points for different movements (Methods). To allow for comparison between different networks that have different neural spaces, we normalized these distances between movements by the distances between adjacent time points along the same latent trajectory, which should be comparable across networks. If the organization within neural space is congruent to the organization in motor output, we would expect the distances in neural space to be proportional to the distances in motor output, and this was indeed the case (Figure 4C).

**Figure 4:**
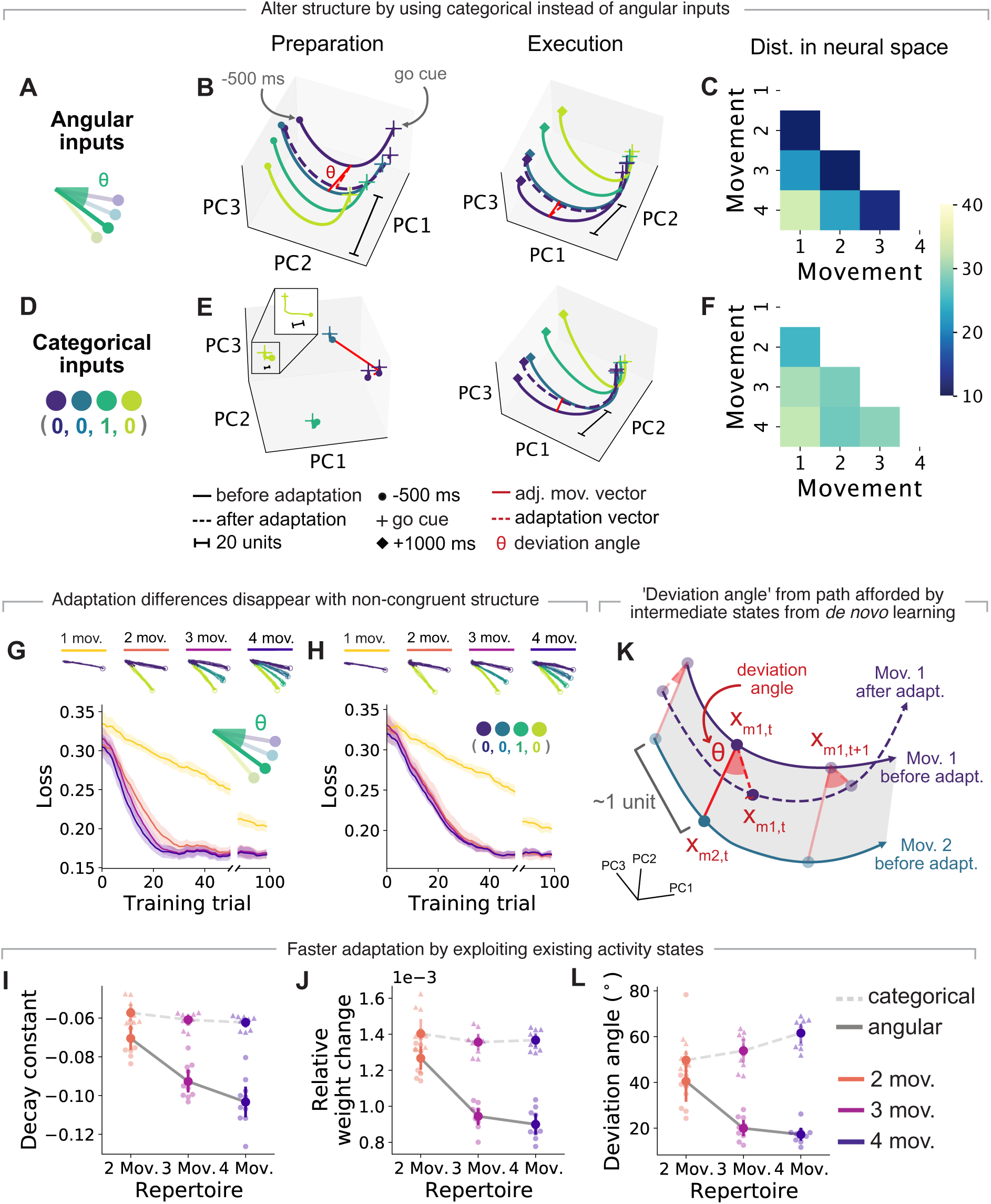
The structure in neural space of multi-movement networks is responsible for their patterns in adaptation. Networks were given either angular (**A**) or categorical (**D**) inputs, and after training on repertoires of one to four movements, then had to adapt to a counterclockwise VR perturbation of 10°. **B**. Latent activity for an example network with angular inputs trained on four movements during preparation (500 ms before go cue) and execution (1000 ms after go cue). Each trace corresponds to the trial-averaged activity for each movement projected on neural manifold computed before adaptation. Solid lines, activity before adaptation; dotted lines, activity after adaptation. **C**. Normalized median Euclidean distances between latent trajectories during preparation and execution for different movements for the network in Panel B. **E-F**. Same as Panels B-C but for an example network with categorical inputs. **G**. Motor output following skill-learning for networks with angular inputs. Bottom: Loss during adaptation training. Traces and shaded surfaces, smoothed mean and 95% confidence intervals across networks of different seeds. **H**. Same as (**G**) but for networks with categorical inputs. **I**. Decay constants for exponential curves fitted to the loss curves in Panel G and Panel H. Circles and error bars, means and 95% confidence intervals with bootstrapping. **J**. Relative weight changes during adaptation. Circles and error bars, means of the median changes across all weights for each seed and 95% confidence intervals with bootstrapping. **K**. We computed a ‘deviation angle’ between the ‘adjacent movement vector’ (red solid line in Panel B, E) and the ‘adaptation vector’ (red dotted line in Panel B, E). **L**. Circles and error bars, means of the median deviation angles across all trials and timesteps for each seed and 95% confidence intervals with bootstrapping. Note the large difference in learning speed (Panel I), relative weight changes (Panel J), and deviation angle (Panel L) following adaptation between networks with angular and categorical inputs.

Our results (Figure 4C) indicate that multi-movement networks’ better ability to adapt to perturbations may be related to having congruent structure in activity space. To demonstrate this, we devised a manipulation that altered the structure of the network activity without affecting its baseline motor performance after *de novo* learning. Thus far, we have used continuous angular inputs that specified the direction to the targets (Figure 4A). We found that we could enforce a different structure by using ‘one-hot encoded’ binary vectors having no angular information (Figure 4D, Methods). With these categorical inputs, the latent dynamics during preparation for different movements no longer had the same organization as the motor output (Figure 4E), becoming less congruent with the organization of the motor output (Figure 4F). This was the case even if the motor output was equally accurate across network classes (Figure 4G,H top) after the initial *de novo* learning phase.

Having these two classes of networks with different degrees of congruency in the structure in their neural activity with respect to the motor output allowed us to directly examine our prediction that greater congruence would aid adaptation. Indeed, while the performance for networks with angular and categorical inputs was comparable following adaptation (Figure 4G,H bottom), adaptation was in general faster for networks with more congruence (angular input networks) than networks with less congruence (categorical input networks) (Figure 4I), suggesting adaptation was easier (compare Figure 4G and H). Within the angular input networks, adaptation was also faster for those with larger movement repertoires (Figure 4G), but this trend was absent for networks with categorical inputs (Figure 4H). Lastly, single-movement networks lacking structure in neural space were unaffected by the difference in input encoding (Figure 4G,H). These results confirm that congruence between the structure of the latent trajectories in neural space and the structure in the motor output is important to facilitate adaptation.

After establishing that structure in neural space shapes motor adaptation, we sought to understand what property of the networks with angular inputs allowed models with greater repertoires to adapt faster. We had previously shown that the structure of multi-movement networks organizes the dynamics and allows them to produce intermediate movements that they have not been trained on (Figure 2G,I; Figure S4B,C). Thus, by having been trained to generate a broader range of motor outputs, the more defined structure of larger repertoire networks in neural space may allow them to produce more intermediate activity states. These intermediate states may facilitate adaptation by avoiding the need to learn new activity patterns. When modeling learning using RNN models, connectivity must be altered in order to change the activity patterns that can be produced ^3, 26^. This implies that if large repertoire networks do exploit intermediate states provided by additional movements, they would require smaller adaptive weight changes than smaller repertoire networks. This was indeed the case, but notably it only occurred in the more congruently structured networks with angular inputs (Figure 4J).

While this confirms that networks with larger learned repertoires require smaller changes for adaptation, it is unclear how intermediate states contribute to these changes. To explore the role of these intermediate states, we examined how the network activity evolved during adaptation. If we assume that intermediate states exist between the latent trajectories for each movement, we would expect the latent trajectories to move along these states during adaptation, in the direction of other latent trajectories. To measure this, we defined a ‘deviation angle’ that quantifies how changes in the trajectories during adaptation deviate from the path afforded by the existing potential intermediate states created during *de novo* learning (Figure 4K; for validation of this metric on motor cortical recordings from monkeys performing the same VR task ^6^, see Figure S6). If the network is using existing intermediate states, we would expect these angles to be small. Indeed, the angular encoding networks that had more congruent structure than the categorical networks also had smaller deviation angles (Figure 4L), with the differences paralleling those seen in the adaptation speed (loss curve decay constants) and relative weight changes (Figure 4I,J).

In summary, how latent activity is structured in neural space has a large impact on motor adaptation. Critically, the existing structure determines the organization of the latent dynamics and allows for intermediate states between learned movements, thereby facilitating VR adaptation only if this structure is congruent to the motor output structure. Without any structure to guide adaptation, single-movement networks adapted the most slowly to our VR perturbation (Figure 5A). Without congruent structure, multi-movement networks with categorical inputs (Figure 5B) adapted more slowly than those with angular inputs (Figure 5C). Finally, with more movements to provide more intermediate states across the structure, networks with the largest repertoires adapted the fastest (Figure 5D).

**Figure 5:**
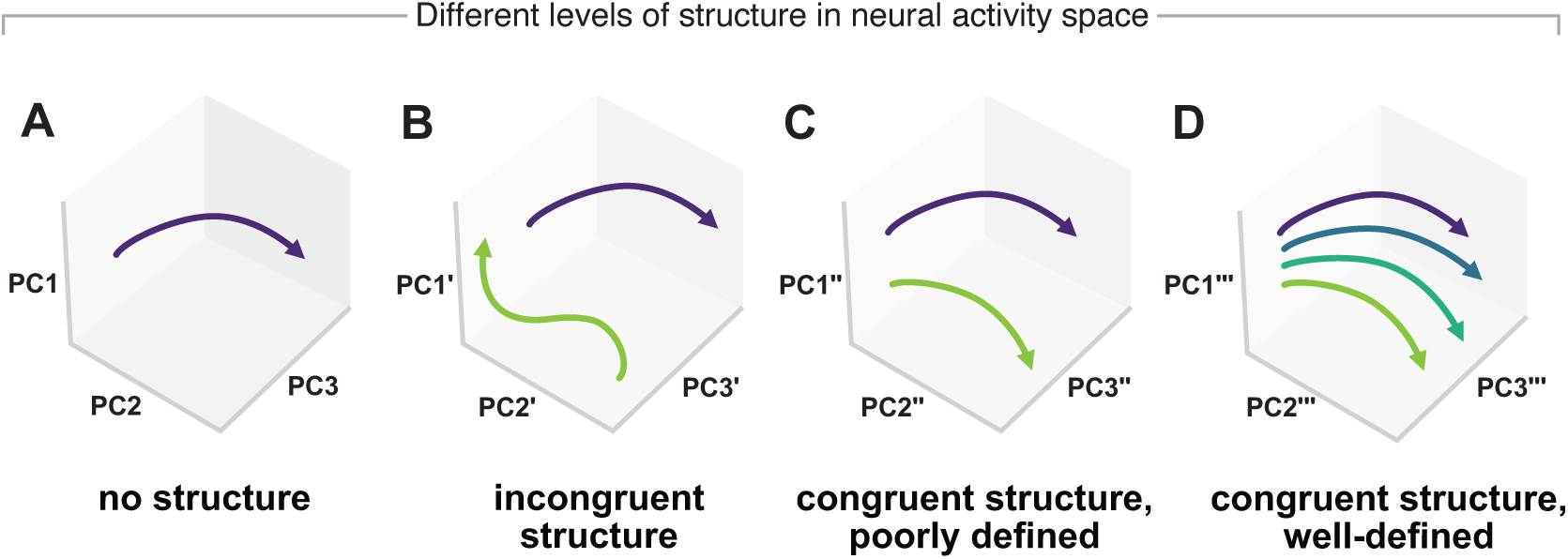
Neural spaces can have different levels of structure. Neural structure is defined as the organization of the latent dynamics in neural space. Networks can have no structure (**A**, e.g. single-movement networks). Networks can have neural structure that is incongruent (**B**, e.g. two-movement networks with categorical inputs) or congruent (**C**, e.g. two-movement networks with angular inputs) to the structure of the motor output. Additional movements can provide more intermediate states to better define the structure (**D**, e.g. four-movement networks with angular inputs)–similar to how additional columns can provide additional support across a building’s framework.

### Structure in neural space can facilitate or impede adaptation

While we found that different degrees of structure in neural space affect adaptation, until now we have only examined adaptation to a VR perturbation. Networks with larger repertoires adapted more quickly only if they had angular inputs. In this case, their latent dynamics were organized by the angular direction of the movements (Figure 4B), allowing them to adapt more quickly to a perturbation that required angular changes in the motor output (Figure 4G). Thus, we hypothesised that the structure across the inputs, the neural space, and the perturbation all need to be congruent for adaptation to be facilitated.

To test this hypothesis directly, we examined how networks adapted to a different type of perturbation that was congruent with the organization of categorical inputs but not with that of angular inputs. In this ‘re-association’ perturbation (Figure 6A), the cues and targets were rearranged such that the network needed to re-associate learned reaches to different known target cues ^26^. Since this perturbation requires adaptation to categorical rather than angular changes, we predicted that multi-movement networks with categorical inputs would adapt more easily than would those with angular inputs.

**Figure 6:**
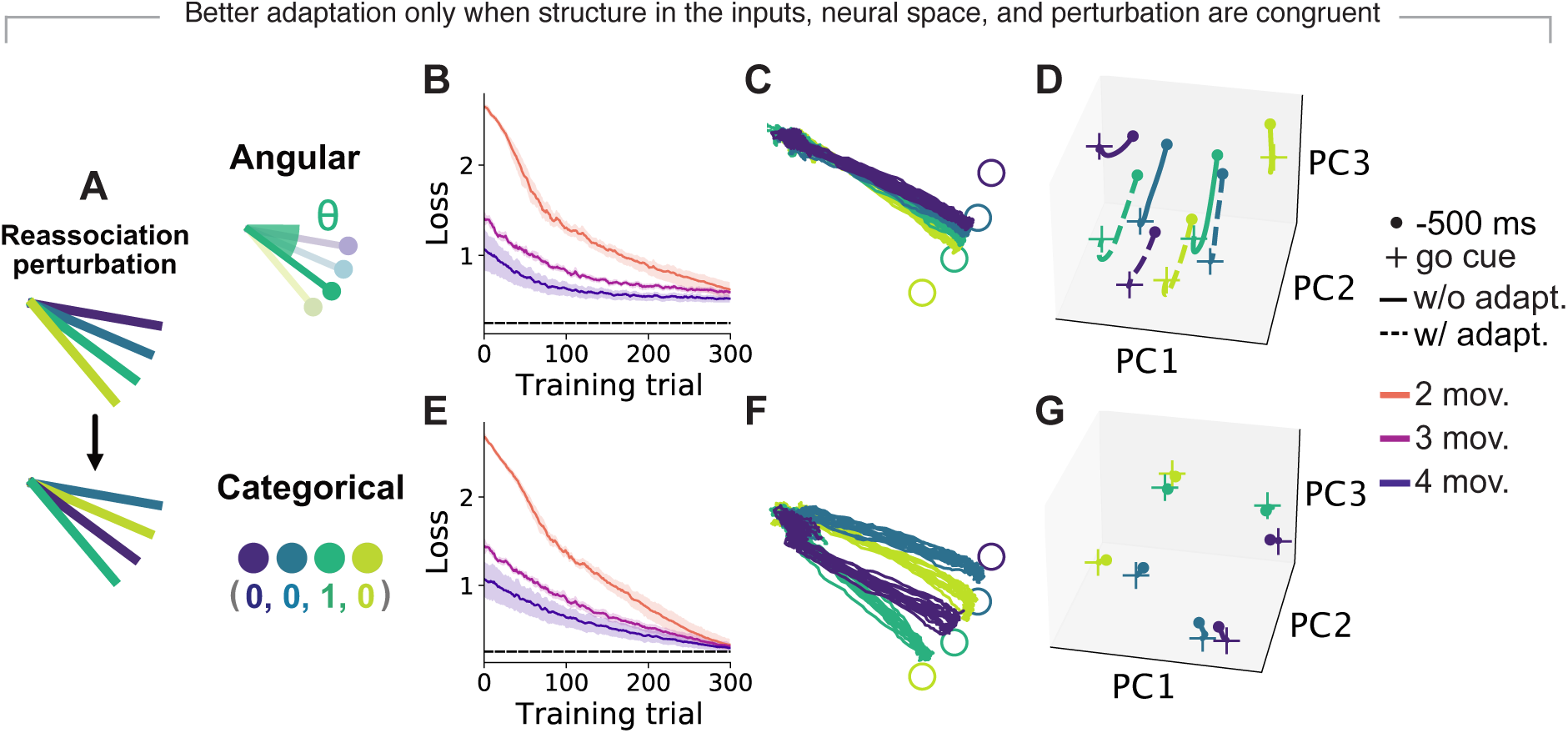
Structure in neural space can facilitate or hinder adaptation. **A**. Networks with angular or categorical inputs adapted to a re-association rather than a visuomotor rotation perturbation. **B**. Loss during adaptation training for networks with angular inputs with varying motor repertoires. Traces and shaded surfaces, smoothed mean and 95% confidence intervals across networks of different seeds. **C**. Motor output following adaptation training. **D**. Latent trajectories during preparation (see 4A) for one example seed. **E-G**. Same as Panels B–D but for networks with categorical inputs. Note that in this case networks with angular encoded could not adapt to the perturbation whereas networks with categorical inputs did.

Indeed, networks with categorical inputs adapted to the re-association perturbation (Figure 6B,C), whereas networks with angular inputs were unable to adapt (Figure 6 E,F) despite comparable performance following *de novo* learning (Figure 4G,H top, Figure S7). For both classes of networks, the latent trajectories maintained their general structure (Figure 6D,G), suggesting that the networks adapted by reusing activity patterns in the existing intermediate states rather than by exploring new ones. However, for networks with angular inputs, neighboring latent trajectories interfered with one another during adaptation such that the adapted motor output became overlapped (Figure 6C,D). Thus, the existing structure in neural space now harmed adaptation instead of facilitating it. Together, these results suggest a fundamental relationship between the structure of neural activity following *de novo* learning and the ability to adapt to a subsequent perturbation: the underlying structure can facilitate adaptation if it is congruent to the perturbation, but it can also hinder adaptation if the perturbation is incongruent. Having well-defined structure in activity space also hindered adaptation to other types of perturbations that required greater changes in the motor output (Figure S8). In conclusion, structure in the neural space can shape adaptation by facilitating adaptation under small changes or interfering with adaptation under larger changes.

## Discussion

Adapting to an external perturbation requires the generation of new activity patterns whose availability is likely shaped through long-term motor skill learning. Motor repertoires from different learning experiences thus provide different baselines for adaptive activity to evolve. Here, we examined how learning different repertoires shapes the underlying structure in neural space and consequently impacts motor adaptation using RNNs. We hypothesized that having a larger repertoire of motor skills could facilitate adaptation since more activity patterns would be readily available. Indeed, we found that networks with larger repertoires could adapt to perturbations more quickly. However, to our surprise, this was true only if (1) the structure of the inputs, the neural space, and the perturbation are all congruent; and (2) small changes in network weights are sufficient. These results suggest that adaptation is affected by both the repertoire of existing activity patterns and the organisation of these patterns in neural space.

### Relation to previous work

Previous work showed that, during a carefully designed BCI adaptation experiment, short-term adaptation could be achieved by re-associating existing neural activity patterns, such that the overall repertoire of neural activity patterns remains the same following adaptation ^27^. Other adaptation studies observed that neural activity was shifted following adaptation ^4, 6^. Notably, however, this shift only occurred under force field perturbation, and not under VR perturbations ^4, 6^, suggesting that our modeling results may fall under the former re-association regime. Indeed, all multi-movement networks readily had the ability to produce the activity patterns necessary for adaptation following *de novo* learning: they were able to adapt to perturbations during adaptation training even with frozen recurrent weights (Figure S7E,F), showing that recurrent weight changes that produce new activity patterns were not necessary. Within this regime, more structured networks still adapted more easily, suggesting that the underlying structure determines how easily activity patterns can be deployed during adaptation. This adds an additional layer of complexity to studies investigating adaptation within the existing neural manifold ^15, 19, 40^.

Furthermore, we uncovered a relationship between input structure, neural structure, and perturbation structure that could lend insight to previous findings. Work on ‘structural learning’ showed that participants adapted more easily to novel perturbations that have the same structure as perturbations from prior experience ^35, 41^. They argued based on behavioural data that knowing this structure facilitates learning by facilitating exploration of a previously acquired low-dimensional ‘task-related’ space. Here, we showed that structure of external cues during learning can shape how the latent trajectories are structured (Figure 4, Figure 6). These latent trajectories define a low-dimensional neural manifold along which further adaptation evolves, providing a potential neural substrate for those behavioural observations.

### Experimental predictions

We can make several experimental predictions based on our results. First, while it is experimentally difficult to examine long-term learning on the timescales we are interested in, we may be able to test our predictions in long training sessions that employ repeated movements. In Verstynen and Sabes ^42^, the authors showed that participants had more varied reach angles towards a given target when a series of recently performed reaches had more variance. These experience-dependent changes in variance mirror those we saw in our simulations (Figure S2), where networks that learned more movements had greater variance and less precision in the motor output (Figure 2E). These similarities suggest that we could perhaps use longer training sessions to study some effects of long-term learning on short-term adaptation. Thus, using a similar experimental setup as Verstynen and Sabes ^42^, we predict that there would be a trade-off between motor output precision and robustness to neural noise ^39^ when more movements are learned through repetition, as shown by our model (Figure 2E,F).

Second, while there were some differences between multi-movement networks of different sizes, the greatest differences were between single and multi-movement networks (Figure 2). Multi-movement networks with the same distribution of learned movements had similar ease in adaptation (Figure 3B,C; Figure S4D,E), since networks were able to produce activity for intermediate movements within the distribution. Thus, we predict that participants would adapt more quickly if they learn a larger distribution of movements, and learning fine-tuned movements within the distribution would provide smaller benefits.

Third, the structure of the inputs shaped how the networks adapted under different perturbations since it changed the underlying structure in neural space (Figure 4, Figure 6). By comparing networks and monkeys trained on the center-out reach task, we saw that the deviation angles in the monkey neural activity were more similar to those for networks with angular inputs (Figure S6). Classically, the center-out reach task has been performed by showing participants the position of the reach target (e.g. Refs. 6,43). Visually, this is perhaps more similar to the angular inputs since it specifies the angular location of the target. To further examine the potential impact of the structure of the inputs on learning, discrete inputs could be used instead by cuing targets with different shapes or colors rather than the explicit target location. Under different types of cues, we would predict different patterns of adaptation under different kinds of perturbations such as a VR or the re-association perturbation we have studied.

### Model limitations and future work

To assess how different motor repertoires affect adaptation, we trained the networks in two stages: *de novo* learning of multiple movements followed by adaptation on a single movement. This set-up makes the networks vulnerable to catastrophic forgetting, since the networks may forget the other movements that are not being trained on during adaptation. Training with FORCE learning ^44^ was especially susceptible to catastrophic forgetting: following adaptation, networks were largely unable to produce output for the other movements (Figure S5). In contrast, training with stochastic gradient descent was largely able to overcome catastrophic forgetting (Figure S5) without overwriting the initial network (Figure S9). Consequently, we decided to use stochastic gradient descent throughout our simulations since it was more behaviorally relevant.

By directly relating the network model activity to motor output, we aimed to model population activity in the motor cortex, which is the main cortical area that projects to the spinal cord to create movement ^45^. While the model has similarities to experimental recordings in monkey motor cortex (Figure S6), it has not been explicitly fitted to neural data (e.g. Refs. 30,46–48). Thus, our model is largely region-agnostic, and it is still unclear where these neural changes due to motor learning and adaptation may occur. Motor learning seems to be associated with activity changes across cortical regions such as the premotor cortex ^5, 49^, primary motor cortex ^6, 50, 51^, parietal cortex ^52, 53^, as well as other structures such as the cerebellum ^54–56^ and perhaps even basal ganglia ^57, 58^. How these different regions interact to affect skill learning and adaptation is an ongoing area of study, and future work could use modular and area-specific networks that are constrained by neural data to tease apart their contributions ^46–48^.

Future work can also examine more complex motor skills. Here, we have focused on simple movements adapted from the center-out reach task since it allowed us to compare the motor output and latent dynamics to experimental data. We can expand on this work to examine more complex and realistic repertoires that include different behaviors like grasping and squeezing, along with more complex modeling of arm kinematics ^59^. Different actions have been shown to occupy different parts of neural state space ^12, 14^, so different combinations of behaviors may alter the underlying manifold and affect subsequent adaptation to perturbations on any given behavior.

Finally, a recent computational model has identified context as a critical factor to unify many aspects of motor learning ^60^. Since our results show that external cues can create structure in neural space (Figure 4), different contexts may similarly be represented by different structures, allowing learning to switch between contexts.

## Conclusion

We have shown that *de novo* skill learning shapes adaptation by creating structure in neural space. This structure is critically modulated by the properties of external inputs during learning, to the extent that two sets of networks that know the same set of movements equally well can exhibit opposite trends when adapting to the same perturbation. Knowing a larger repertoire of movements often facilitates adaptation, but only under certain conditions. First, this holds true only if small changes are needed, demonstrating trade-offs in skill acquisition. Second, the perturbation must be aligned with the structure of the underlying latent dynamics, further highlighting how external cues during learning can shape structure in neural space and subsequent adaptation. While we have examined this formation of structure in the context of motor learning, similar structural constraints may arise in other systems, shaping not only motor but also cognitive processes.

## Methods

### Task

We trained recurrent neural networks to perform a standard center-out reach task, which is commonly used in experimental settings to examine motor control. We modified existing experimental data for reaches using the task design of Ref. 6. In the experiment, a monkey controlled a cursor on a computer screen using a 2-D manipulandum. The cursor starts in the middle of a circle with a radius of 8 cm and it must reach to possible targets spaced around the circle. The monkey is shown which target to move to at a target cue, but they must delay movement until a later go cue. To understand how existing skill sets affect motor adaptation, we created four skillsets of different sizes, ranging from one to four reach movements to one to four targets, respectively. We adapted experimental reaches of one monkey, Monkey M, from Ref. 6 to create the target reaches for each movement by rotating the experimental reaches to different targets that we defined. In our case, the targets were equally spaced around an arc of the circle. Unless specified otherwise, the arc spanned from −10° to −50°. Networks were trained on one of these four skillsets. By using different numbers of movements in each skill set, we were able to examine how a network that knows more movements may adapt differently than a network that knows less. Each trial lasted 4.0*s*: the target and go cues were randomly selected for each trial, with the target cues between 1.0-2.5*s* and the go cues between 2.5-3.0*s*.

To examine network performance without trial-to-trial movement variability, which was inherent in experimental reaches, we also created skillsets with synthetic reaches. These synthetic reaches had varied go and target cues, as in the experiments, but the target position profiles of each reach was the same across trials. Each synthetic reach lasted 1.0*s* and the position profile was defined by a sigmoid for each time *t* for the length of the reach *l* = 8*cm*:

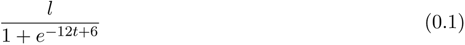

To assess motor adaptation, we examined how networks adapted to a visuomotor rotation (VR), a common perturbation used in experimental settings. To simulate VR, we rotated the output position of the network counterclockwise by *θ_r_* during adaptation trials. We used rotations with *θ_r_* = 10°, 30°, or 60°. We also examined how networks adapted to visuomotor re-associations where the target cues and targets are rearranged, such that the network must reach to a different known target given a known target cue.

### Neural network model

#### Network architecture

The model dynamics were given by:

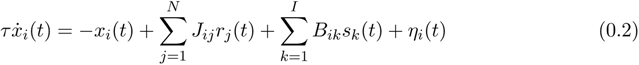

where *x_i_* is the hidden state of the *i*th unit and *r_i_* is the corresponding firing rate following *tanh* activation of *x_i_*. The network has *N* = 300 units and *I* inputs. The time constant *τ* = 0.05*s*, the integration time step *dt* = 0.01*s*, and the noise *η* is randomly sampled from the Gaussian distribution 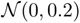 for each time step. The initial states *x_t_*_=0_ are sampled from the uniform distribution 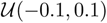. The network is fully recurrently connected, with the recurrent weights *J_ij_* initially sampled from the Gaussian distribution 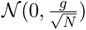, where *g* = 1.2. The time-dependent stimulus inputs *s* (specified below) are fed into the network, with input weights *B* initially sampled from the uniform distribution 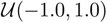.

Two types of sustained inputs *s*were used in our networks. For the angular inputs, *s* is 3-D and consists of a 1-D fixation signal and a 2-D target signal (2 *cos θ^target^,* 2 *sinθ^target^*) that specifies the reaching direction *θ^target^* of the target. For the categorical inputs, *s* is 5-D and consists of a 1-D fixation signal and a 4-D one-hot encoded target signal with the same amplitude (e.g. (0,0,2,0)) that does not provide information about the target’s angular direction. For both types of inputs, the fixation signal starts at 2 and goes to 0 at the go cue and the target signal remains at 0 until the task cue.

The networks were trained to produce 2-D outputs *p* corresponding to *x* and *y* positions of reach trajectories, and they are read-out via the linear mapping:

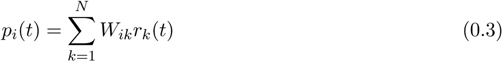

where the output weights *W* are sampled from the uniform distribution *U* (*−*1.0, 1.0). During VR adaptation trials, *p* is rotated counterclockwise according to a perturbation angle *θ_r_*.

### Training the model

Networks were optimized to generate positions of reach trajectories modified from Ref. 6. The training and testing datasets were created by pooling successful trials during baseline epochs across all experimental sessions for Monkey M (2208 trials: 90% training, 10% test). The experimental data was modified for each repertoire (see Task), and equal numbers of trials for each reach direction were included for each repertoire.

Unless otherwise specified, networks had to learn their input weights *B* and recurrent weights *J* while the output weights *W* remained fixed. To model motor skill learning, we initially trained the networks on skillsets with one to four movements, using the Adam optimizer with an initial learning rate *l* = 10*^−^*^4^, first moment estimates decay rate *β*_1_ = 0.9, second moment estimates decay rate *β*_2_ = 0.999, and epsilon *ɛ* = 1*e −* 8. Then, to model motor adaptation, we trained the pre-trained networks to either counteract a VR or re-associate targets and target cues, using stochastic gradient descent with a fixed learning rate *l* = 5*^−^*^3^, unless otherwise specified. We used a faster learning rate during adaptation to model faster short-term learning compared to long-term skill learning. To assess how existing skillsets may shape adaptation on a given movement, networks were only trained to counteract the VR on the one target that all skillsets share (i.e. the −10° reach), such that performance was comparable across networks with different skillsets. Initial training was implemented with 750 training trials and a batch size *B* = 64. Adaptation training was implemented with either 100 training trials for VR perturbations or 300 training trials for reassociation perturbations, and a batch size *B* = 64. All training configurations were performed on 10 different networks initialized from different random seeds. To examine adaptation under an alternate learning algorithm, we also trained the networks to counter the VR perturbation using FORCE learning ^44^ with a learning rate of 100.

The loss *L* was the mean squared error between the 2-D output and target positions over each time step *t*, with the total number of time steps *T* = 400. The first 50 time steps were not included to allow network dynamics to relax:

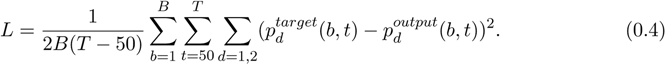

To produce dynamics that align more closely to experimentally estimated dynamics ^28, 29^, we added L2 regularization terms for the activity rates and network weights in the overall loss function *L_R_* used for optimization:

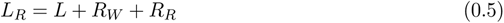

where

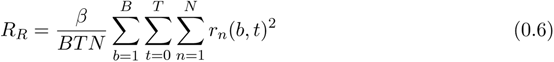

And

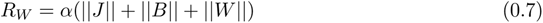

where *β* = 0.5 and *α* = 0.001. Note that the loss recorded in the main text was the loss *L* before regularization, and it was smoothed with a backward moving average of five trials. We clipped the gradient norm at 0.2 before applying the optimization step.

## Data Analysis

Analyses on the neural activity were examined for both the preparation and execution epochs of the movement, taken as 500 ms before and 1000 ms after the go cue, respectively.

To assess how latent dynamics change during motor learning and adaptation, we examined the neural activity space of the networks. In the activity space of a population of *n* neurons, each point denotes the state of the neural population, and each axis corresponds to the firing rate of a specific neuron. To obtain smooth firing rates through time, we applied a Gaussian kernel (std = 50 ms) to the activity rates from the networks. We identified a lower *m*-dimensional neural manifold in the activity space by applying PCA to the smoothed firing rates of the neurons. PCA finds orthogonal basis vectors (principal components or PCs) that maximally captures the variance in the population activity. PCA finds *n* PCs for an *n*-dimensional space, but we sorted the PCs by their corresponding eigenvalues to get the *k*-leading PCs, or neural modes, that capture the majority of the variance. Here, we used *k* = 10, which captured more than 80% of the variance in our network activity. We projected the original smoothed firing rates onto the neural modes to get the latent dynamics of the networks.

To measure the variability in activity and motor output, we aligned trials by the go cue and measured the variance across trials for the same reach at corresponding time points. To compare the variance in latent dynamics across different neural spaces, we first normalized the latent dynamics by the median distances between trial-averaged time points within the neural space before calculating the variance. Variance in unit activity and latent dynamics was calculated during both preparation and movement, while variance in output position was calculated during movement. Reach angles were calculated based on the mean angle for the entire reach during movement, and the variance was calculated across trials. To determine how patterns in variance in the motor output can differ from those in the activity, we examined the variance in unit activity in the output-potent and output-null subspaces ^38^. Unit activity was directly related to the motor output through the read-out weights *W* of the networks. The output-potent dimensions were then the row space of *W* while the output-null dimensions were the null space of *W*. We projected the unit activity onto these dimensions to get the activity in the respective subspaces.

To assess how the underlying neural manifold changes during motor adaptation, we visualized these changes by projecting both the unit activity before and after adaptation onto the neural modes of the manifold before adaptation (Figure 4B, E). We quantified the manifold overlap before and after adaptation training as specified in Ref. 32, adapted from Ref. 61. To find the manifold overlap between the manifolds of two network activities *A*_1_ and *A*_2_, we first find the covariance matrix *C*_1_ of *A*_1_ and project it onto its neural manifold identified through PCA. We then find the covariance matrix *C*_2_ of *A*_2_ and project it onto the neural manifold of *A*_1_. To quantify the variance explained by these projections, we divide the trace of these projections by the trace of the corresponding covariance matrices:

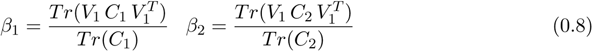

where *V*_1_ are the first 10 principal components resulting from PCA on *A*_1_. Here, *β*_1_ is the variance in *A*_1_ that can be explained by the neural manifold for *A*_1_ while *β*_2_ is the variance in *A*_2_ that can be explained by the neural manifold for *A*_1_. We then calculate the manifold overlap as the ratio *β*_2_/*β*_1_.

Changes in the neural manifold are driven by changes in synaptic connectivity, so we also measured the relative weight change *dW* before and after adaptation training:

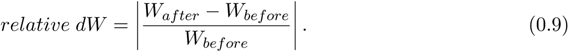

To examine the direction of these changes in the manifold relative to the initial shape of the manifold, we defined a metric called the ‘deviation angle’. First, we define ‘adjacent movement vectors’ *v_adj_* between corresponding time points of the trial-averaged latent trajectories of the first movement **x***_m_*_1_ and its adjacent movement **x***_m_*_2_ (the second movement) before adaptation:

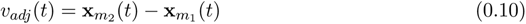

These vectors quantify the general shift from one movement to the next in neural space and approximate the shape of the manifold between the adjacent movements. Then, we perform a similar computation for the first movement before (**x***_m_*_1_) and after (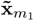) adaptation to get the ‘adaptation vector’ *v_adp_* that quantifies the general shift during adaptation:

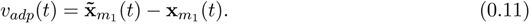

We define the ‘deviation angle’ as the angle between these two vectors, which measures how changes in adaptation deviate from the path afforded by the existing scaffolding before adaptation.

To measure the degree of structure in the neural space, we quantified the median distance *D* between the trial-averaged latent dynamics **x**_1_ and **x**_2_ for each pair of movements over all corresponding time points *t*:

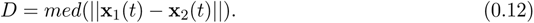

To compare these distances between movements across different neural spaces, we normalized by the median distances between time points within each movement, pooled across all movements *m*:

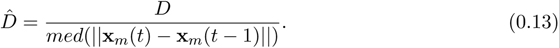

Neural spaces with more structure should have similar distances in neural space between latent trajectories for movements that have similar distances in motor output.

### Experimental comparison

To verify that the networks produced realistic latent dynamics, we trained the networks on the original 8-target center-out reach task defined in Ref. 6 (see Task). The target trajectories for training and testing were based on reach trajectories during successful trials pooled from all sessions for Monkey M (see Training the model). Following training, we compared the simulated activity to experimental activity recorded from motor cortex during one baseline session for Monkey M ^6^. We pre-processed the experimental recordings by removing units with trial-averaged firing rates less than 5 Hz and applying a Gaussian kernel (std = 50 ms) for the binned square-root transformed firings of each unit (bin size= 30 ms). We then substracted the cross-condition mean. To compare the latent dynamics (see Data Analysis), we used Canonical Correlation Analysis (CCA), which finds new directions (canonical correlations or CCs) in the neural manifold that maximize the pairwise correlations between two data sets when they are projected on these directions ^62^. Canonical correlation values that are associated with these CCs range from 0 to 1, with 1 being complete correlation.

To verify the ‘deviation angle’ metric, we trained the networks to counteract a 30° VR perturbation on all 8 targets and compared the deviation angles to those found in two monkeys performing the same adaptation (3 sessions for Monkey C, 3 sessions for Monkey M). Deviation angles were found for the latent dynamics corresponding to reaches for all 8 targets. We also calculated deviation angles for shuffled targets and time points as a control for the networks. To compare the deviation angles with the extent of adaptation, we calculated error curves for the monkeys based on the angular error of their reaches in the first 150*ms* and calculated decay constants for exponential curves fitted to these error curves, as we did for the loss curves during training for the networks.

## Data availability

The data that support the findings in this study are available from the corresponding authors upon reasonable request.

## Code availability

All code to reproduce the main simulation results will be made freely available upon publication on GitHub.

## Author contributions

J.C.C., J.A.G. and C.C. devised the project. M.G.P. and L.E.M. provided the monkey datasets. J.C.C. ran simulations, analysed data and generated figures. J.C.C., C.C. and J.A.G. interpreted the data. J.C.C., C.C. and J.A.G. wrote the manuscript. All authors discussed and edited the manuscript. J.A.G. and C.C. jointly supervised the work.

## Competing Interests

J.A.G. receives funding from Meta Platform Technologies, LLC.

## Acknowledgements

J.C.C. received funding from the Wellcome Trust (grant 108908/Z/15/Z). M.G.P. received funding from the Fonds de recherche du Québec Santé (grant chercheurs-boursiers en intelligence artificielle J1). L.E.M. received funding from the NIH National Institute of Neurological Disorders and Stroke (NS053603 and NS074044). J.A.G. received funding from the EPSRC (EP/T020970/1) and the European Research Council (ERC-2020-StG-949660). C.C received funding from the BBSRC (BB/N013956/1 and BB/N019008/1), the EPSRC (EP/R035806/1), the Wellcome Trust (200790/Z/16/Z), and Simons Foundation (564408). The funders had no role in study design, data collection and analysis, decision to publish, or preparation of the manuscript.

## Supplementary Figures

**Supplementary Figure S1:**
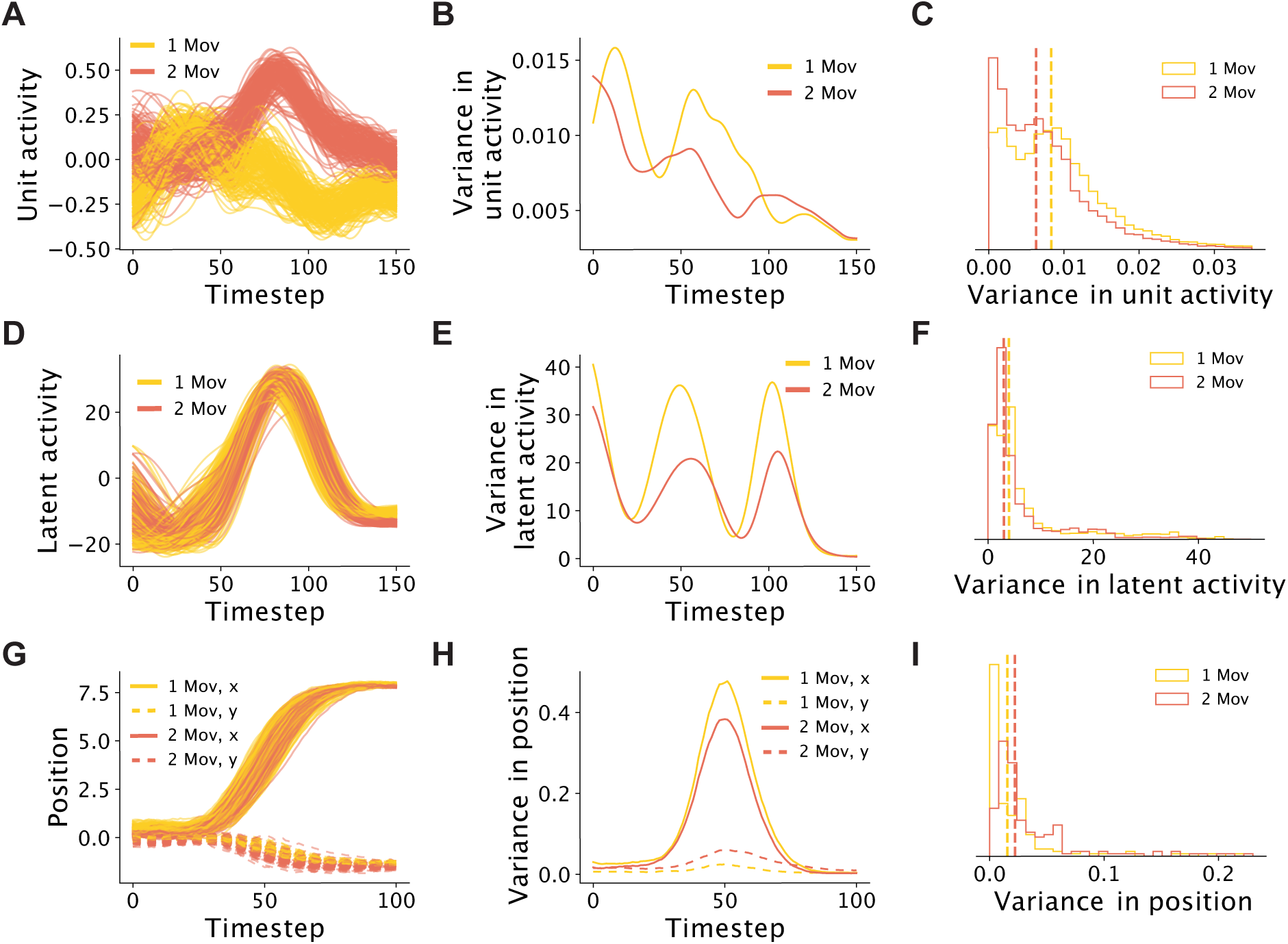
Networks that can generate multiple movements produce more constrained neural dynamics. This figure presents additional data for Figure 2B-D, and compares different variables across networks with different repertoires producing the same common movement. **A**. Unit activity for one example unit for networks trained on one (yellow) or two (orange) movements. Networks had the same random seed. Traces, different trials. **B**. Variance in unit activity, calculated per timestep across all trials for example unit in Panel A. **C**. Distributions for variance in unit activity, pooled across all units for one example seed. Dashed lines, median. **D-F**. Same as Panels A-C but for the 2nd dimension of the latent dynamics in Panels D-E and for all dimensions in Panel F. **G-I**. Same as Panels A-C but for the motor output. In Panels G-H: Solid line, position along the *x* axis; dotted line, position along the *y* axis.

**Supplementary Figure S2:**
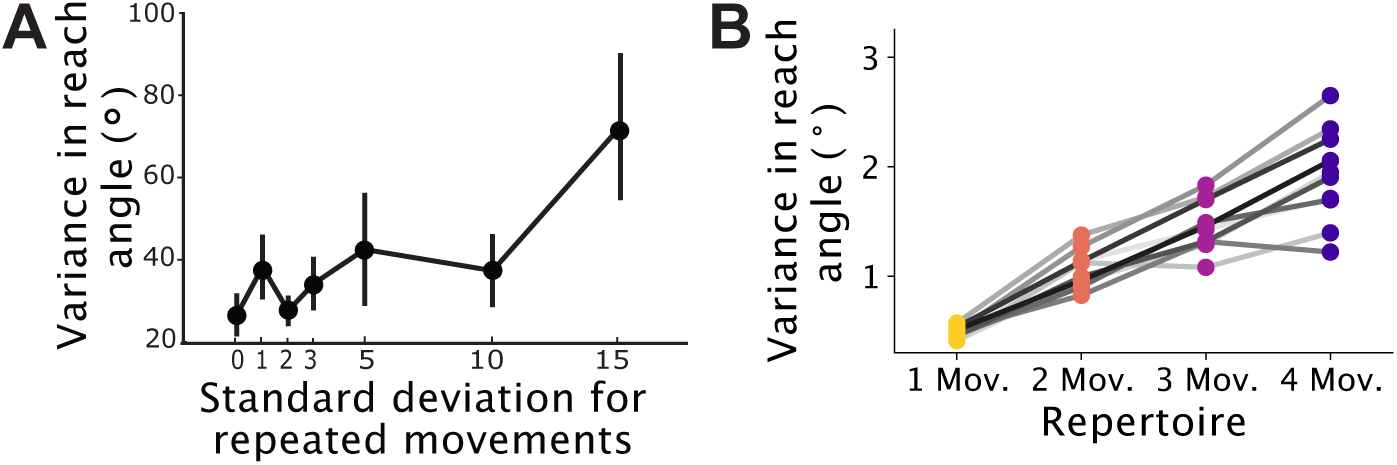
Networks reproduce experimental results from Ref. 42. **A**. This study examined the variance in reach angles to a given target following repeated movements sampled from a normal distribution around the target with different standard deviations. Larger variance of known movements was correlated with larger variance in reach angle. Figure modified from Fig. 2A from Ref. 42. **B**. Networks initially trained on different repertoires were tested on one shared movement. Networks trained on larger repertoires had a larger variance of known movements, and this was also correlated with larger variance in reach angle.

**Supplementary Figure S3:**
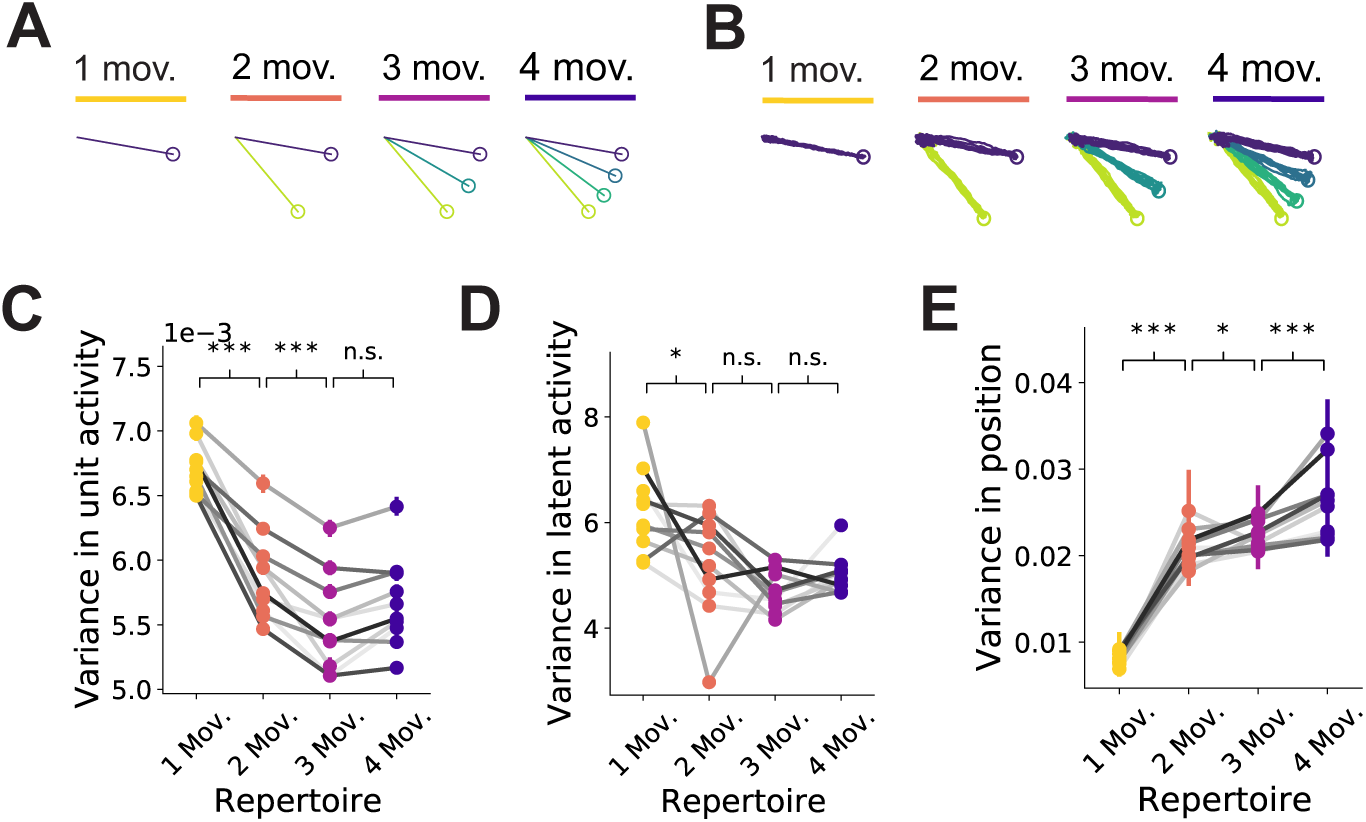
Constrained dynamics are not a byproduct of variability in the monkey movements they were trained on. Networks were trained to produce motor output (**B**) based on simulated (**A**) rather than actual “hand trajectories”. **C-E**. Same as Fig.2C-D but for networks trained on simulated hand trajectories. Note that the patterns remained the same as when trained on actual hand trajectories.

**Supplementary Figure S4:**
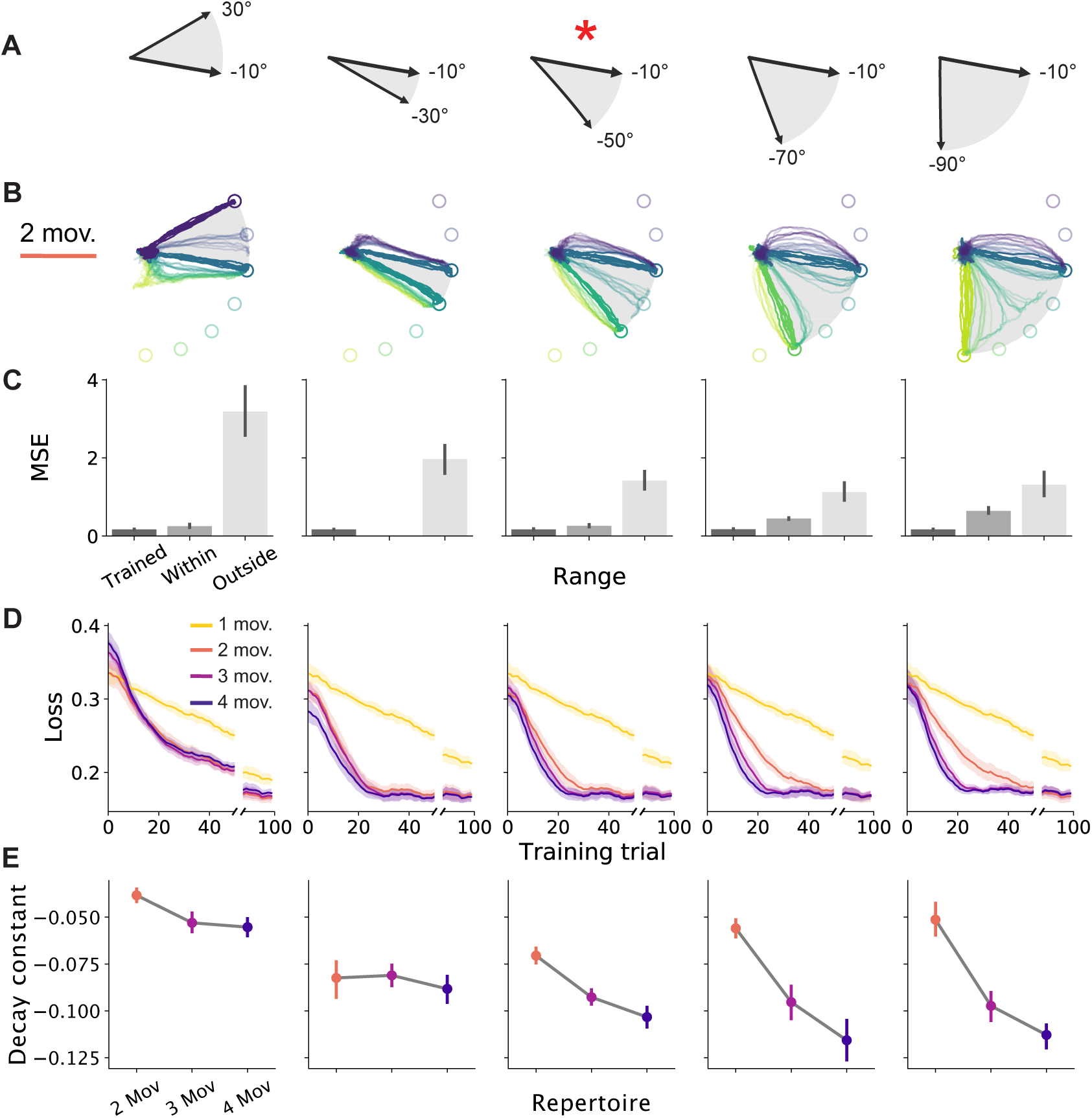
Networks can generalize and adapt to perturbations that require movements within a learned range. **A**. Networks were trained on repertoires with movements that spanned different ranges (top). The range −10° to −50°, denoted by an red asterisk, was used for all simulations in the main text. **B-C**. Networks trained on two-movement repertoires in their respective ranges (i.e. a movement to −10° and a movement to −50° for the −10° to −50° range) were tested on target cues for movements equally spaced between 30° and −90° to assess whether they could generalize to movements they were not trained on. The target cues were chosen such that networks were tested on movements they knew (‘Trained’), movements that were within the range of known movements (‘Within’), and movements that were outside the range (‘Outside’). **B**. Motor output for the target cues for each movement for a sample network. Colors, target cues; circles, targets for each movement. Colors for target cues that the networks have not been trained on have lower opacity. Grey backgrounds denote the range of known movements. **C**. Mean-squared error between the network output and target positions for previously ‘Trained’ movements, and movements ‘Within’ or ‘Outside’ the range of known movements. Note that MSE was lower for ‘Within’ movements, showing that the networks can generalize. **D**. Loss during adaptation training with counterclockwise VR perturbations of 10°. Traces and shaded areas, mean and 95% confidence interval across networks of different seeds. **E**. Decay constants for exponential curves fitted to the loss curves in Panel D. Circles and error bars, mean and 95% confidence interval with bootstrapping.

**Supplementary Figure S5:**
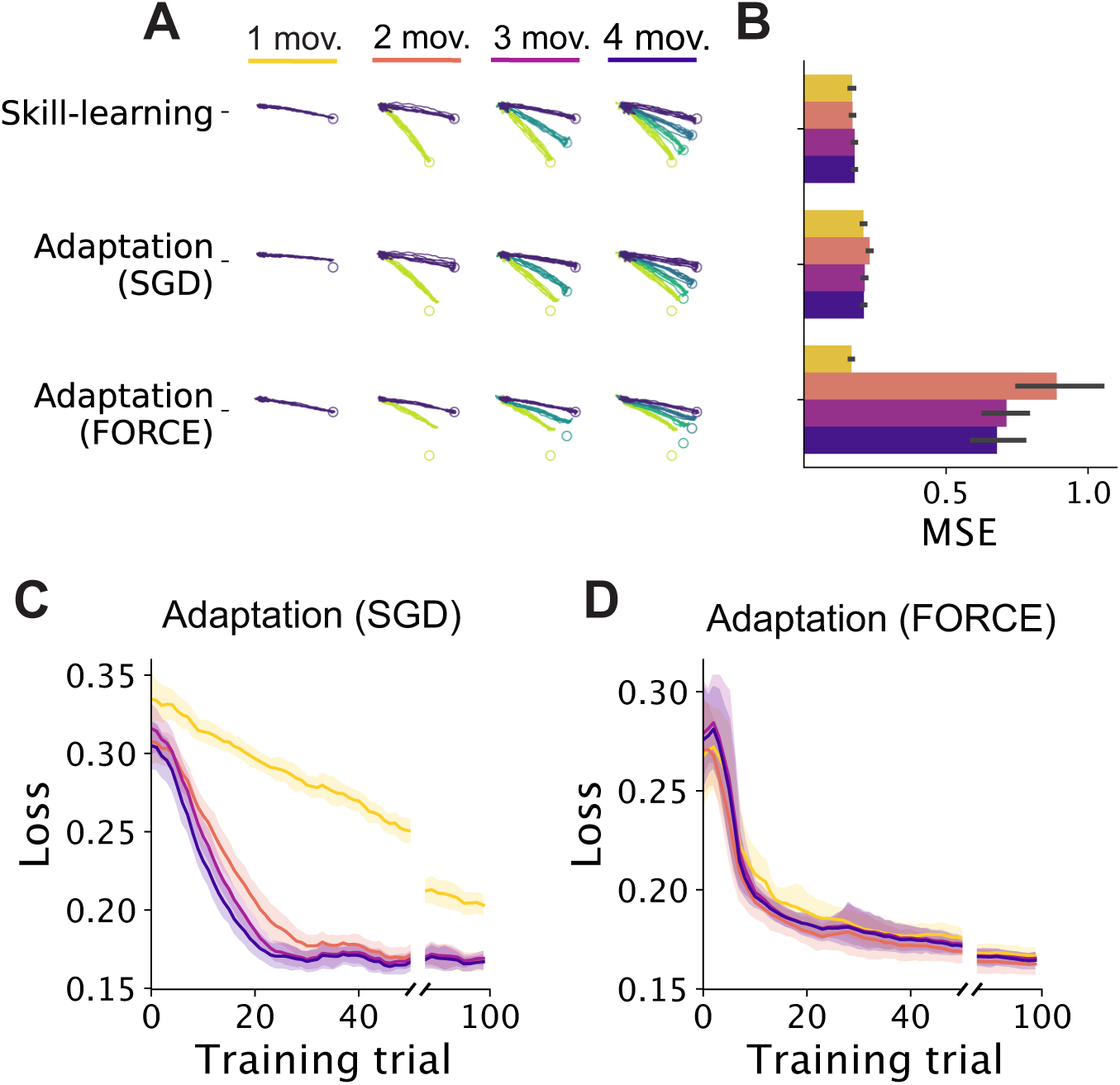
Adaptation training with stochastic gradient descent (SGD) is better at overcoming catastrophic forgetting than FORCE. Following skill learning, networks were trained to adapt to counterclockwise VR perturbations of 10° on one movement, using either SGD or FORCE. Motor output (**A**) and MSE (**B**) for all movements in known repertoires following adaptation training. Note that networks trained with FORCE had worse ‘catastrophic forgetting’ (that is, forgetting of previously trained tasks when learning new tasks) of other movements that were not perturbed during adaptation training. Loss during adaptation training with SGD (**C**) and FORCE (**D**). Traces and shaded areas, mean and 95% confidence intervals across networks of different seeds.

**Supplementary Figure S6:**
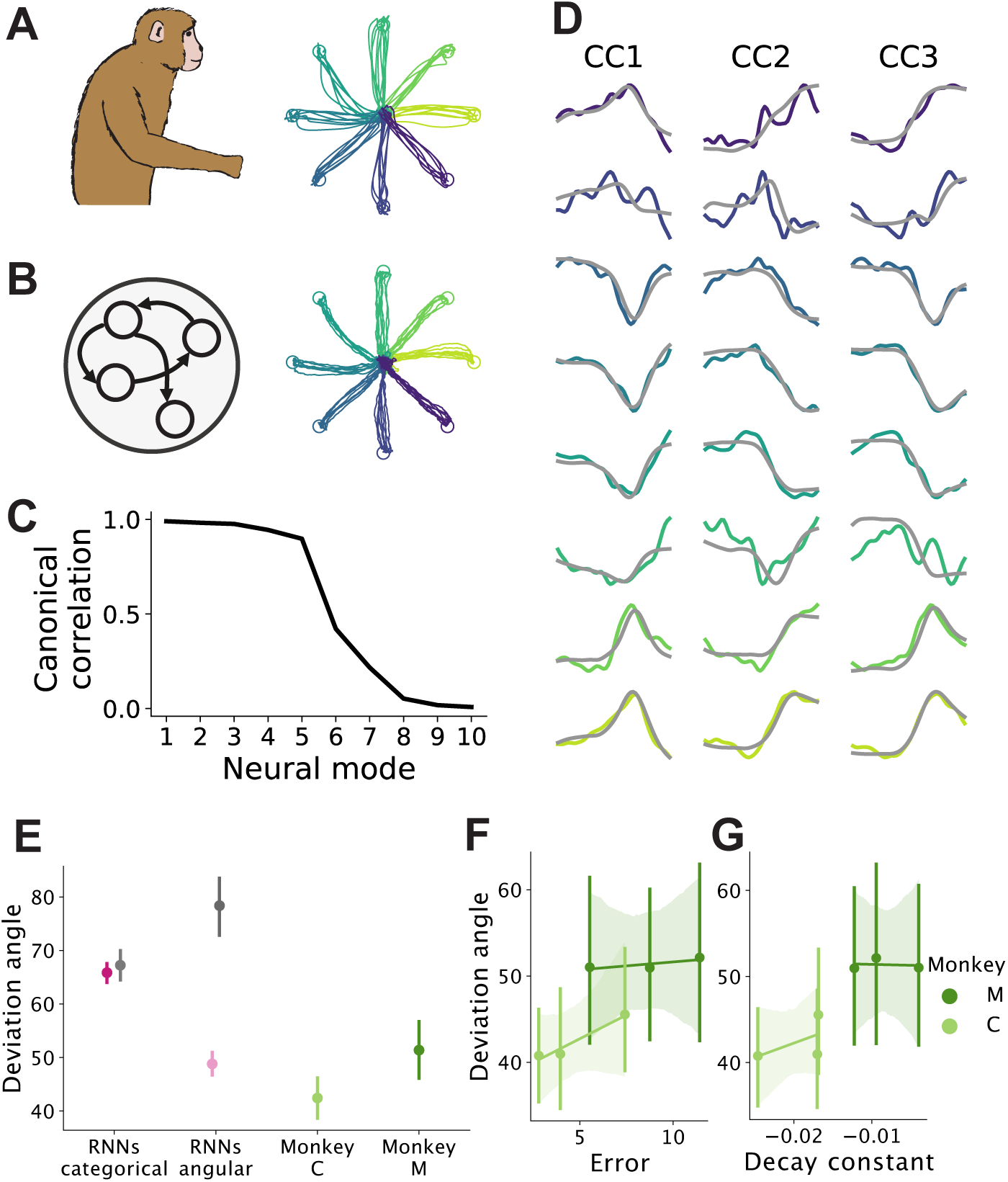
Recurrent neural networks produce realistic neural dynamics. **A**. Hand trajectories produced by a monkey performing an eight-target center-out reaching task. Data from Ref. 6 (pooled from all sessions for Monkey M, see Methods). **B**. Simulated motor output when RNNs were trained on the same center-out reaching task with angular inputs. **C** Canonical correlation values between experimental and simulated latent dynamics. **D**. Projections of the experimental (colored) and simulated (grey) latent dynamics onto the first three axes identified through canonical correlation analysis. **E-G**. RNNs were trained on the center-out reaching task with either one-hot encoded (‘categorical’) or angular inputs, and compared to two monkeys trained on the same task (Monkey C and Monkey M). Data from Ref. 6. **E**. The ‘deviation angle’ (Figure 4K) was calculated and pooled for the latent activity of all targets before and after adaptation to a visuomotor rotation. Circles and error bars, median and 95% confidence intervals with bootstrapping. Grey, control with shuffled targets and time points. **F**. Deviation angles compared to the angular error of reaches for the monkeys for each session. **G**. Deviation angles compared to the decay constants fit to learning curves based on the angular reach errors. Note that Monkey C had smaller deviation angles and learned faster (more negative decay constants).

**Supplementary Figure S7:**
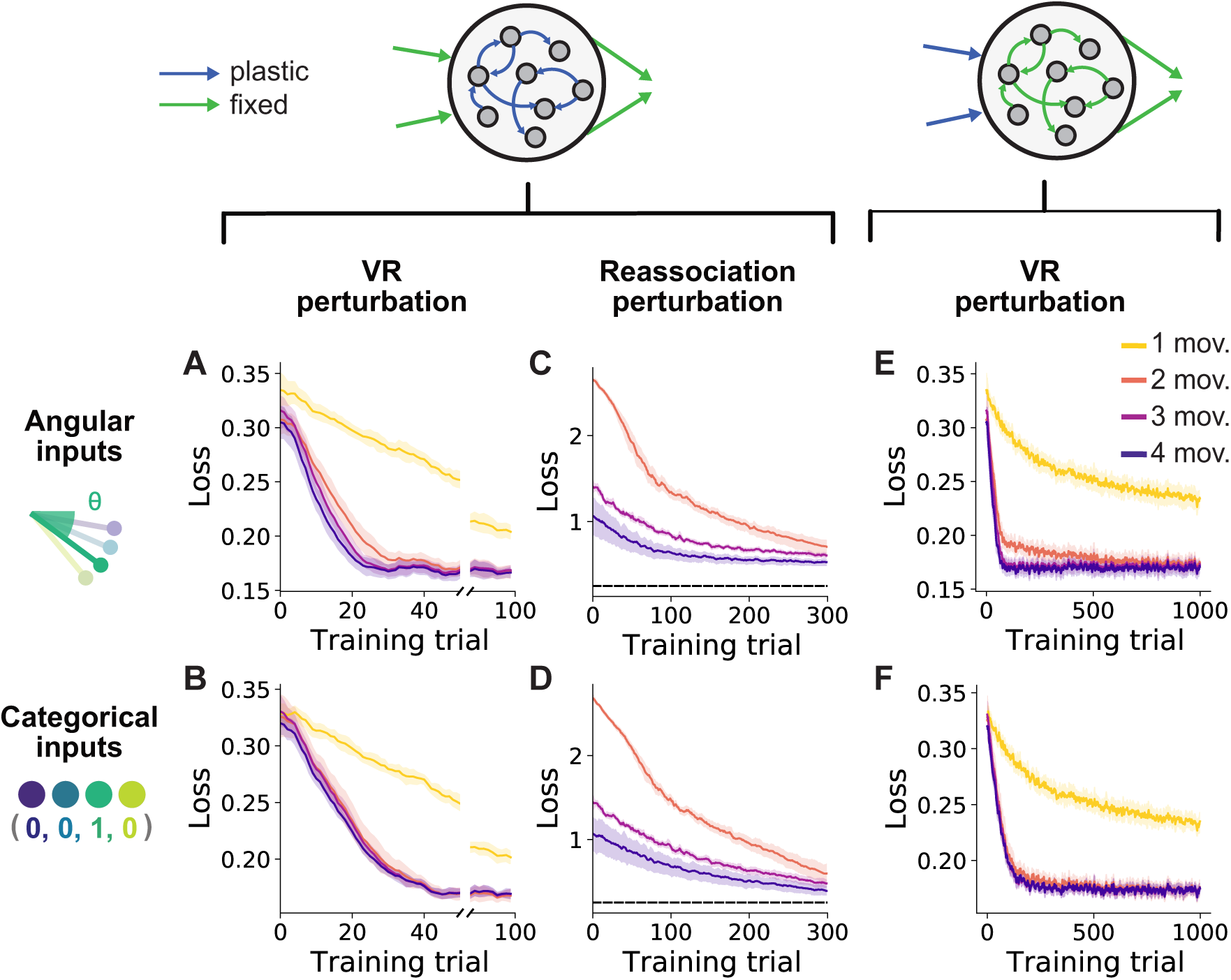
Adaptation with fixed input or recurrent weights. Loss during adaptation training with 10° counterclockwise VR perturbations with fixed input weights for networks with angular (**A**) and categorical (**B**) inputs (for 100 training trials). Traces and shaded areas, smoothed mean and 95% confidence intervals across networks of different seeds. **C, D**. Same as Panel A, B but for reassociation perturbations (for 300 training trials). Note that the results from Figure 6A,D held when the input weights were frozen such that the networks could not simply rely on changing the input weights to counteract re-association. **E, F**. Same as Panels A, B but with fixed recurrent weights (for 1000 training trials). Note that the patterns remain the same as in the main text, where networks were trained with plastic input and recurrent weights, although at longer timescales.

**Supplementary Figure S8:**
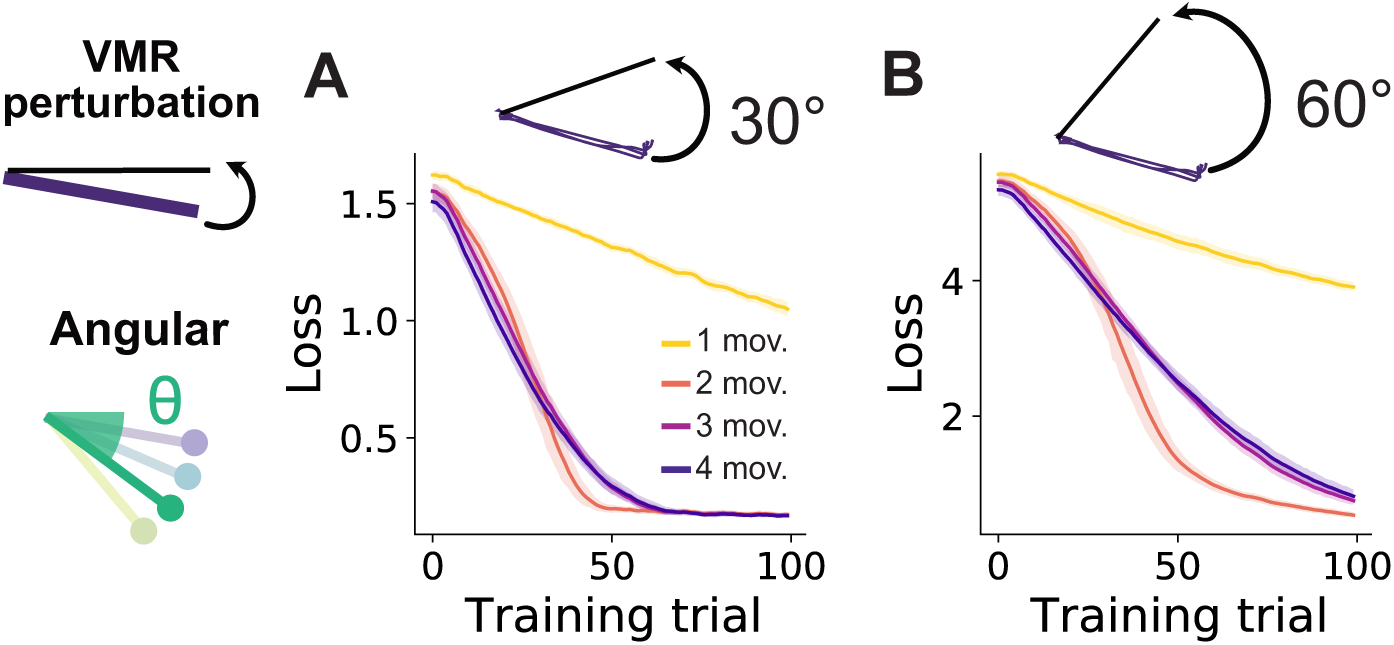
Structure in neural space hinder adaptation when larger changes are required. Networks were given angular inputs and adapted to ‘more challenging’ VR perturbations than the 10°rotations examined thus far. **A**. Loss during adaptation training for a VR of 30°. **B**. Loss during adaptation training for a VR of 60°. For both perturbations, smaller multi-movement networks adapted more quickly than larger multi-movement networks, contrary to previous results under smaller perturbations of 10° (Figure 3D). Note that this effect occurred when the movement needed to counter the adaptation was both within (for the 30° VR) and outside the range (for the 60° VR) of the known movements.

**Supplementary Figure S9:**
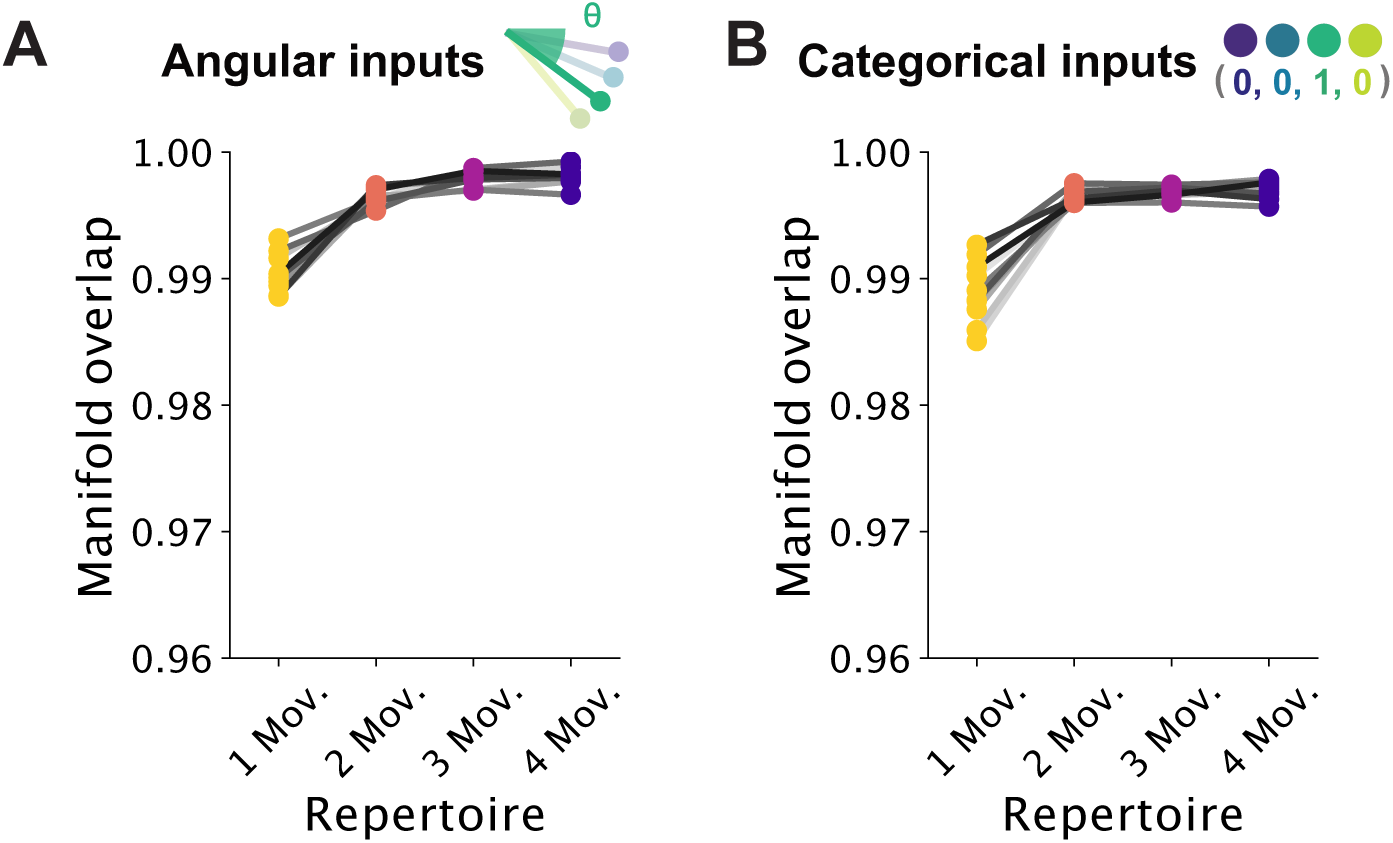
Adaptation to VR perturbation is within-manifold. Manifold overlap (see Methods) for network activity between skill learning and adaptation to a VR perturbation of 10° for networks with angular (**A**) and categorical (**B**) inputs. Line colors denote different random seeds.

## Notes

### Summary of Updates

Author names updated

